# Adaptive repulsion of long-term memory representations is triggered by event similarity

**DOI:** 10.1101/2020.01.14.900381

**Authors:** Avi J. H. Chanales, Alexandra G. Tremblay-McGaw, Brice A. Kuhl

**Affiliations:** Department of Psychology, New York University, 6 Washington Pl., New York, New York 10003, USA; Department of Psychology, 1227 University of Oregon, Eugene, Oregon 97403, USA; Institute of Neuroscience, University of Oregon, Eugene, Oregon 97403, USA

## Abstract

We tested whether similarity between events triggers an adaptive repulsion of long-term memories. Subjects completed an associative learning task in which objects were paired with faces. Critically, the objects consisted of pairs that were identical except for their color values, which were parametrically varied in order to manipulate interference. Performance on associative memory tests confirmed that color similarity robustly influenced interference. Separate tests of color memory showed that high similarity triggered a repulsion of long-term memories, wherein remembered colors were biased away from colors of competing objects. This repulsion effect was replicated across three experiments. In a fourth experiment, the repulsion effect was fully eliminated when task demands promoted integration, instead of discrimination, of similar memories. Finally, we show that repulsion of color memory was highly adaptive: greater repulsion was associated with less memory interference. These findings reveal that similarity between events triggers adaptive distortions in how events are remembered.

**SUPPLEMENTARY INFORMATION:** Three supplementary figures are included.

## INTRODUCTION

When episodic memories are similar, this creates interference and, ultimately, can lead to forgetting. Thus, an important challenge for the memory system is to resolve interference so that forgetting is minimized. The hippocampus is thought to play a critical role in resolving interference by pattern separating memory representations (Bakker, Kirwan, Miller, & Stark, 2008; Colgin, Moser, & Moser, 2008; Yassa & Stark, 2011). Recently, several neuroimaging studies have found that hippocampal activity patterns associated with highly similar memories systematically ‘move apart’ from each other, suggesting that interference triggers a *repulsion* of memory representations (Ballard, Wagner, & McClure, 2019; Chanales, Oza, Favila, & Kuhl, 2017; Dimsdale-Zucker, Ritchey, Ekstrom, Yonelinas, & Ranganath, 2018; Favila, Chanales, & Kuhl, 2016; Hulbert & Norman, 2015; Schlichting, Mumford, & Preston, 2015). However, these fMRI findings raise an important question: does a similar repulsion also occur with respect to how the specific *features* of competing events are remembered?

Here, we report a series of behavioral experiments—directly inspired by evidence of hippocampal repulsion—that test whether competition triggers repulsion of feature values associated with competing long-term memories. We had two central predictions. First, feature repulsion should be *competition dependent*—repulsion should be more likely to occur when memories are highly similar to each other (Chanales et al., 2017; Schapiro, Kustner, & Turk-Browne, 2012). Second, feature repulsion should be adaptive—repulsion should be associated with a reduction in memory interference (Favila et al., 2016; Hulbert & Norman, 2015).

To test these ideas, we conducted four experiments, each using a similar long-term, associative memory paradigm. In the paradigm, subjects repeatedly studied and were tested on object-face associations. Although each face was associated with a unique object, we created competition by including object pairs that were identical except for their color values (e.g., a blue backpack and a purple backpack; Figure 1A). Moreover, we parametrically manipulated the color distance between these object pairs to precisely control the degree of competition. In addition to measuring subjects’ associative memory (i.e., which face was paired with which object), we also probed subjects’ memory for the color of each object, providing a critical measure of whether subjects exaggerated the feature distance between competing objects. To the extent that competition triggers memory repulsion, we expected repulsion in color memory to specifically occur when color similarity was high. To the extent that repulsion is adaptive, we expected greater repulsion to be associated with less associative memory interference.

**Figure 1:**
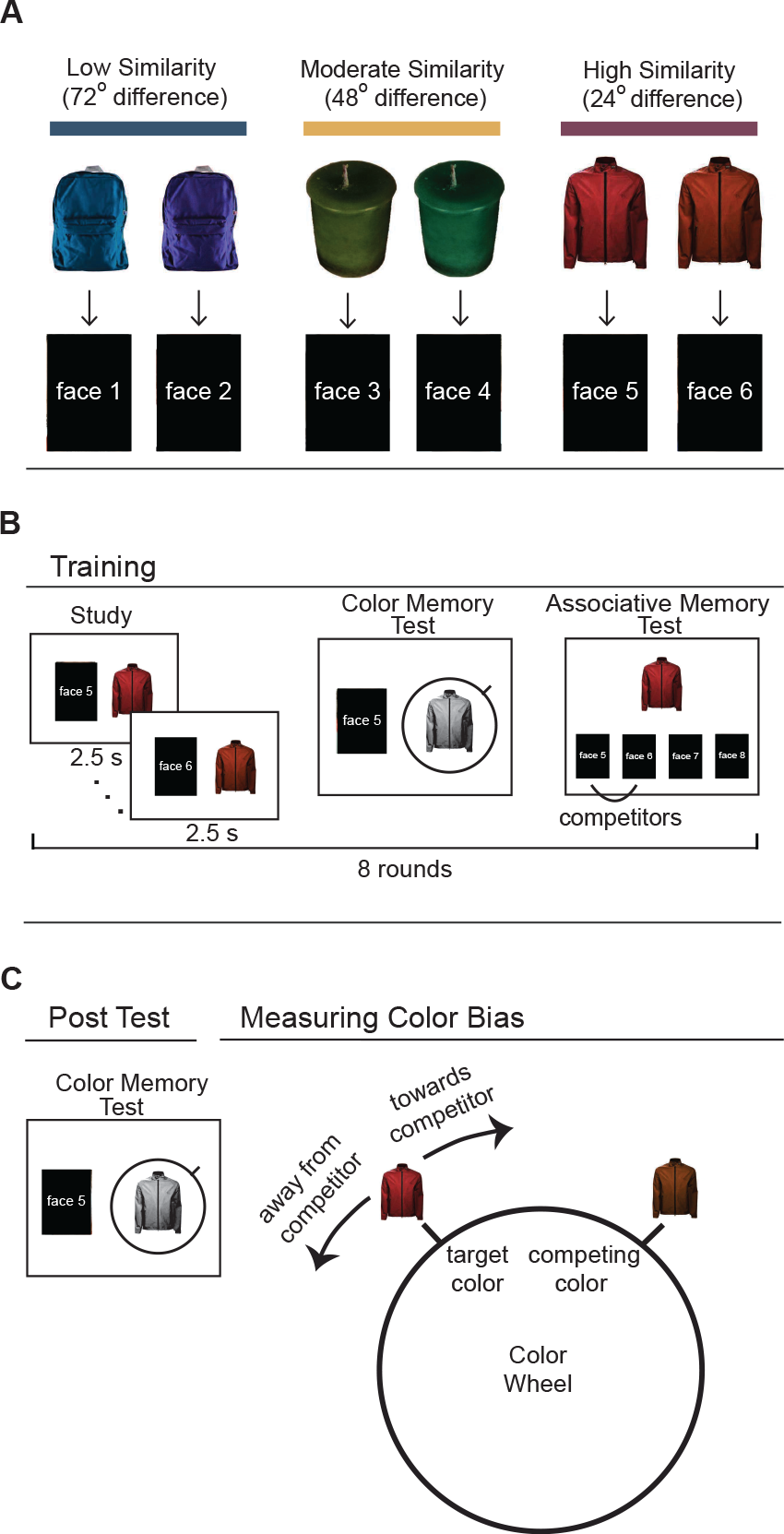
Experimental Design. **(A)** In each of four experiments, subjects learned object-face associations that contained pairs of competing objects (object images that were identical except for their color values). The similarity (color distance) between competing objects was parametrically manipulated within and across experiments. In Experiments 1, 2, and 4, there were three similarity conditions: high (24 degrees apart), moderate (48 degrees), and low (72 degrees). In Experiment 3, the conditions were: moderate (48 degrees), high (24 degrees), and ultra (6 degrees). Note: Actual faces are not shown here per biorxiv policy. **(B)** Each experiment began with 8 training rounds. Each training round contained a study, a color memory test, and associative memory test phase. During study (*left panel*), subjects viewed each object-face pair. During color memory tests (*middle panel*), subjects were presented with a face and a greyscale version of the associated object. Using a continuous color wheel, subjects selected (recalled) the color of the object. During associative memory tests (Experiments 1–3 only; *right panel*), an object image was presented and subjects selected the associated face from a set of four options. The four face options always included the correct face (target) and the face that had been paired with the competing object (competitor). Procedures for Experiment 4 are described in **Figure 5** and in Methods. **(C)** For all experiments, after the training rounds subjects completed a Post Test that only probed color memory. The procedure was identical to the color memory tests from the training rounds. The critical performance measure from the Post Test was the percentage of color memory responses that were biased away from the color of the competing object.

## METHODS

### Participants

For all Experiments, target sample sizes were identified in advance, but final sample sizes were determined by exclusion criteria. Because we did not have any way to estimate effect sizes in advance of the first Experiment, we chose a target sample of 40 subjects for Experiment 1. However, due to exclusion criteria, a total of 23 subjects (11 males, 18-32 years old) were included in analyses. Instead of adding additional subjects to increase the sample size for Experiment 1, we conducted a replication study (Experiment 2) with a larger sample: 36 subjects included in analyses (13 male, 18-22 years old). For Experiments 3 and 4, the number of subjects included in analyses were: 38 subjects in Experiment 3 (6 males, 18-34 years old) and 26 subjects in Experiment 4 (1 male, 18-25 years old). The rationale for the smaller sample in Experiment 4 was that pilot data indicated that the critical manipulation in Experiment 4 was quite powerful and consistent across subjects. Exclusion criteria are described below, in Procedures. All subjects were right-handed and reported normal or corrected-to-normal vision. Informed consent was obtained in accordance with procedures approved by the University of Oregon Institutional Review Board.

### Materials

For all experiments, stimuli consisted of 18 object images and 36 face images (all images were 250 x 250 pixels). The object images were selected from a set of images designed to be color-rotated (Brady et al., 2013). Objects were chosen that had no strong inherent associations with particular colors. To alter the color of each object, the hue of an image was rotated through a circular color space ranging from 0–360 degrees. Colors were altered by independently rotating every pixel through an equiluminant circle in L*a*b* space. Face images were pictures of non-famous white males gathered from the internet. For each subject, 6 object images were randomly assigned to each of three color similarity conditions. For Experiments 1, 2, and 4, these conditions were: high similarity (24 degrees), moderate similarity (48 degrees), and low similarity (72 degrees) (Figure 1A). For Experiment 3, these conditions were: ultra similarity (6 degrees), high similarity (24 degrees), and moderate similarity (48 degrees). Each object was then assigned a pair of colors separated by the hue angle degree difference of their respective condition. For Experiments 1, 2, and 4, this was accomplished by randomly selecting 45 colors from the color space, each separated by 8 degrees (45 * 8 = 360 degrees). This set of 45 colors represented the set of possible colors for each subject, but only 36 of these colors were actually used (18 objects * 2 colors per object). For Experiment 3, the ultra similarity condition necessitated a slight modification to the color assignment procedure: 60 colors (instead of 45) were randomly selected from the color space, each separated by 6 degrees (instead of 8). This set of 60 colors represented the set of possible colors for each subject, but, again, only 36 of these colors were used. For all Experiments, the 36 colors were then assigned to objects, according to their similarity condition, without replacement (i.e., each color was only assigned to one object). One constraint on this assignment was that, for each condition, there was an even representation across each third of the color space (1-120, 121-240, 241-360 degrees).

### Procedures

The first part of each experiment was a series of training rounds (8 total) in which subjects learned and were tested on all of the object-face associations (Figure 1B). Specifically, each training round was comprised of a study phase, a color memory test phase, and an associative memory test phase. During the study phase, subjects were shown (encoded) each object-face association. During the color memory test phase, subjects were presented with a face image and a greyscale version of the object that was associated with that face. Using a continuous report scale (Brady, Konkle, Alvarez, & Oliva, 2013), subjects selected (recalled) the color of the object. During the associative test phase (with the exception of Experiment 4), subjects were presented with an object image along with four face images and they attempted to select (retrieve) the face that had been paired with that object. Importantly, the set of four faces included the face that was paired with the object (‘target’), the face that was paired with the competing object (‘competitor’), and two faces that had been paired with different objects (‘non-competitors’). After the 8 training rounds, subjects completed a color memory Post Test which repeatedly tested subjects’ memories for each object’s color (Figure 1C). This Post Test was identical in format to the color memory tests during the training rounds, but served as a critical measure of the ‘end point’ of subjects’ learning.

#### Experiment 1

Experiment 1 consisted of 8 training rounds and two Post Tests. Each training round included study, color memory test, and associative memory test phases (in that order). In each study round, subjects viewed the same 36 face-object associations. For each trial, a face and corresponding object image appeared on a white screen for 2.5 s, with the face image to left of the object. There was a 1 s inter-trial interval during which a blank white screen was presented. Each object-face association was studied once per study round.

On each color memory test trial, a studied face was presented to the left of its paired object and subjects used a color wheel (Brady et al., 2013) to select the remembered color of the object. The object image initially appeared in grayscale; once participants moved the mouse cursor, the object would appear in color. The hue was determined by the angle between the cursor location and the center of the object image. A line marker along a ring surrounding the object image indicated the current hue angle. Once subjects rotated to the desired color they clicked the mouse to finalize their choice. There was no time limit for these responses. The color wheel was randomly rotated across trials so there was no correspondence across trials between spatial position and color.

The associative memory tests probed memory for each object-face association. On each trial, a colored object was presented at the top of the screen and four faces images were presented beneath. Subjects were instructed to select the face that had been studied with the object image. The target face was always presented along with a face that was paired with the competing object. Thus, subjects had to discriminate between the objects’ colors (in memory) in order to select the correct face. The other two face images were two randomly selected faces that had been paired with other, non-competing objects (non-competitor faces). Each face served as a non-competitor foil on exactly 2 trials. Subjects made a selection using a computer mouse with no time limit to respond. They then indicated confidence in their response by clicking either a ‘sure’ or ‘unsure’ button using the mouse (note: these confidence ratings are not considered here). A feedback screen was then presented for 1.25 s; the feedback screen indicated whether the selected face was correct or not and also displayed the correct object-face pair.

Following the 8 training rounds, subjects completed an immediate Post Test (Day 1 Post Test) and then returned the following day (~24 hours later) for a second Post Test (Day 2 Post Test). The Post Tests were identical to the color memory tests in the training rounds except that, in the Post Tests, each object was tested 5 times. The 180 post-test trials were divided into 5 blocks in which each object was tested exactly once. The order of the trials within a block was randomized with the constraint that an object and its competitor were not tested on successive trials. To minimize fatigue, after trial numbers 60 (1/3 of trials competed) and 120 (2/3 of trials completed) a screen prompted subjects to “Take a quick break” and to press a key to continue. The Day 2 Post Test was identical to the Day 1 Post Test except that the order of trials was re-randomized.

#### Experiment 2

Experiment 2 was identical to Experiment 1 except for a few procedural changes. Because of the strict performance-based exclusion criteria in Experiment 1, and the time limit cutoff (1.5 hours), there was a high overall exclusion rate (45.2%) and relatively small final sample of subjects (n = 23). Thus, the goal for Experiment 2 was to replicate the results from Experiment 1, but with a larger final sample (the target was a 50% increase) and lower overall exclusion rate. We retained the same exclusion criteria, but sought to shorten the experiment so as to reduce the number of subjects that failed to complete the experiment in the allotted time (1.5 hours). We opted to shorten the experiment rather than to extend the time limit due to concern for subject fatigue. Subject fatigue was of particular concern given that the most critical data from the entire experiment came from the Post Test (i.e., the last round of the experiment). To shorten the experiment, we reduced the number of color memory test rounds during training so that they only occurred during rounds 1, 3, 5, and 7, and we imposed a time limit of 10 s on the associative memory test trials and color memory test trials (both during the training rounds and the Post Test). The number of Post Test trials excluded because of the time limit in Experiment 2 was very small (mean across subjects = 0.78%; maximum = 3.3%). Additionally, since we observed qualitatively identical results across the Day 1 and Day 2 Post Tests in Experiment 1, we did not include the Day 2 Post Test in Experiment 2.

#### Experiment 3

Experiment 3 was identical in procedure to Experiment 2 except that color memory tests were included in each of the 8 training rounds (as in Experiment 1) and the time limit for the entire experiment was extended to 2 hours. The rationale for reverting to every-round color memory tests was that the magnitude of the repulsion effect in the high similarity condition was somewhat lower in Experiment 2 (M = 54.63%) than in Experiment 1 (M = 60.80%) and we speculated that this difference might be partly attributable to the reduction in the number of color memory tests during training in Experiment 2. The rationale for extending the time limit was to reduce the number of subjects excluded for not completing the experiment in the allotted time.

#### Experiment 4

Experiment 4 differed in procedure from Experiments 1–3 most significantly in that the associative memory test during the training rounds was replaced by an *inference test* that assessed generalization across object-face pairs. Specifically, on each trial in the inference test one of the 36 face images appeared at the top of the screen, with four face images presented beneath. Subjects were instructed to select which of the four face options were associated with the same object category (e.g., “backpack”) as the probe face at the top of the screen. Subjects made their selection using a computer mouse. The set of four face options always included the target face (correct response) and 3 other studied, non-target faces. Note: here, the ‘competitor’ face was the target face. Each face was tested once per round (i.e., 36 trials per round) and each face served as a non-target face option on exactly 3 trials. Feedback was provided on each trial indicating whether the selected face was correct along with the correct face-face pairing displayed on screen for 1 second. Otherwise, all procedures were identical to Experiment 3 (for details of color similarity conditions, see Materials).

#### Exclusion criteria

Across Experiments 1–4, a total of 19, 18, 9, and 6 subjects, respectively, were excluded from analysis. Subjects were excluded from analyses if they failed to complete the experiment in the allotted time (1.5 hours for Experiments 1 & 2 and 2 hours for Experiments 3 & 4). For Experiments 1–4, this resulted in exclusion of 12, 0, 1, and 3 subjects, respectively. Additionally, subjects were excluded if they did not satisfy each of two performance-based criteria. The first performance criterion was that, across the last two rounds of the associative memory test (see Procedures for details), *non-competitor* face images were selected on no more than 2% of trials. Importantly, this exclusion criterion was orthogonal to subjects’ ability to discriminate between similar colors in that it only required that subjects had ‘narrowed down’ the options to either the target or competitor face. This exclusion criterion therefore specifically ensured that subjects had learned that two different faces were paired with a common object category (e.g., “backpack”). Across Experiments 1–4, a total of 6, 15, 9, and 0 subjects, respectively, failed to meet this criterion. The second performance criterion was that the percentage of Post Test trials with reaction times less than 500 ms could not exceed 15%. Given that the Post Test trials required clicking and dragging a cursor along a color wheel, responses that were made in less than 500 ms were considered to be evidence of subjects rushing through the experiment—which was a particular concern given the repetitive and tedious nature of the experiment. Across Experiments 1–4, a total of 5, 3, 1, and 6 subjects, respectively, failed to meet this criterion. Note: some subjects failed to satisfy both of the performance-based criteria (Experiments 1–4: 4, 5, 3, and 0 subjects, respectively). It is important to emphasize that these performance-based exclusion criteria were established in advance, they were orthogonal to our effects of interest (repulsion of color memory), and they were applied uniformly across all experiments.

#### Measuring Color Memory

For color memory tests (during training rounds and Post Test), responses were recorded as integer values (0–359 degrees) reflecting hue angle on the color wheel. Although the color memory tests were identical in procedure during the training rounds and Post Test, for narrative clarity we focus on different measures during each phase. During the training rounds, we focus on *color error* as a general measure of training-related improvement in color memory. Color error was computed as the absolute value of the hue angle difference between a subject’s color response and the true color. During the Post Test, however, because we were critically interested in whether color memory responses exhibited bias, we focused on the *percentage of responses away from the competing object’s color*. To illustrate how this measure was computed, the location of the target object’s color value can be considered as being at 0 degrees on the color wheel and a competing object might, for example, be at 24 degrees. In this scenario, any responses between 181 and 359 degrees would be counted as ‘away from’ the competing color. Notably, the definition of ‘away’ responses would be identical if the competing object color was at 6, 48, or 72 degrees. For each subject and each condition, we computed the percentage of trials that fell in the ‘away’ bin. We used this measure as opposed to mean signed error because mean signed error is highly susceptible to influence from extreme responses whereas the percentage of away responses is not. While we focus on the color error measure during training rounds and the percentage of away responses during Post Test, we also report the percentage of away responses during training rounds in **Supplementary Figure 1** and mean color error during Post Test in **Supplementary Figure 3**.

## RESULTS

### Experiment 1

Across the training rounds in Experiment 1, there were marked reductions in error on the color memory tests (Figure 2A) and increases in accuracy on the associative memory tests (Figure 2B) (see **Supplementary Figures 1** and **2** for additional data from the training rounds). Critically, accuracy on the associative memory tests was strongly influenced by color similarity (Figure 2B). In particular, subjects were much more likely to select the face associated with the competing object (hereinafter, *interference errors*) when color similarity was high (**Supplementary Figure 2**), confirming that our interference manipulation was successful. In order to test for repulsion effects in color memory, we focused on the Post Tests. We predicted that repulsion would specifically occur when competition was high (i.e., the high similarity condition). The critical dependent measure was the percentage of trials in each similarity condition for which subjects reported a color that was ‘*away from*’ the color of the competing object (measures of unsigned color error are reported in **Supplementary Figure 3**). For example, if the target object’s color was located at 0 degrees on the color wheel and the competing object’s color was at 24 degrees, a color response at 350 degrees would be considered ‘away from’ the competing object’s color (Figure 1C and see Methods). We defined a repulsion effect as occurring for a condition if the mean percentage of away responses was greater than 50% (i.e., that most color reports were biased *away from the color of the competing object*).

**Figure 2.**
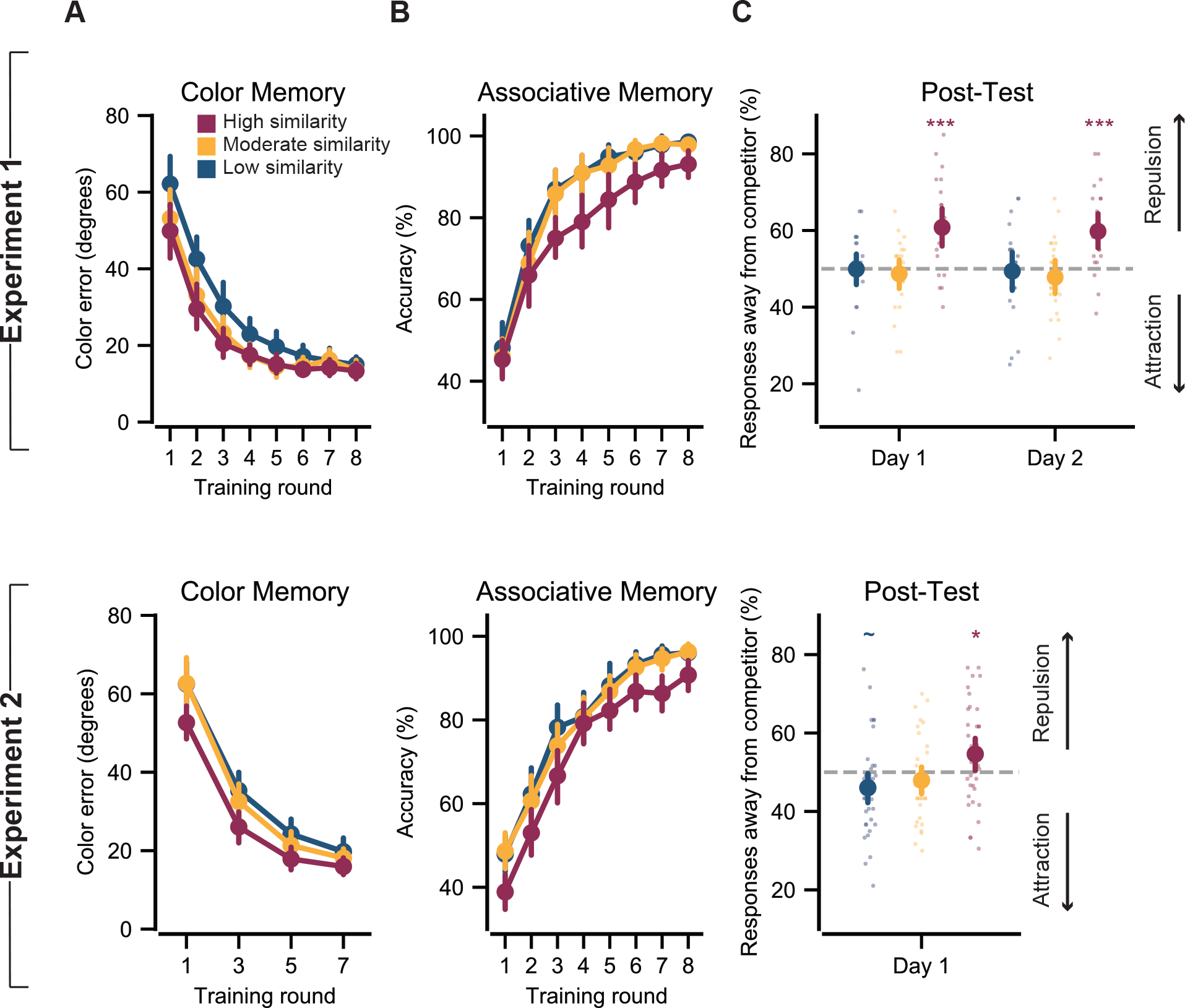
Color similarity induces interference and triggers memory repulsion. Results from Experiment 1 (top row) and Experiment 2 (bottom row). **(A)** Mean color memory error (absolute distance between reported and target color values from the color memory tests) decreased across training rounds (main effect of round, Experiment 1: *F*_1,22_ = 166.2, *P* < 0.0000001, η^2^ = 0.77; Experiment 2: *F*_1,22_ = 166.2, *P* < 0.0000001, η^2^ = 0.83) **(B)** Accuracy on the associative memory tests (percentage of trials for which the target face was selected; chance = 25%) increased across training rounds (main effect of test round, Experiment 1: *F*_1,22_ = 435.4, *P* < 0.0000001, η^2^ = 0.83; Experiment 2: *F*_1,35_ = 690.4, *P* < 0.0000001, η^2^ = 0.85). Accuracy differed across color similarity conditions (main effect of similarity, Experiment 1: *F*_2,44_ = 13.04, *P* = 0.00003, η^2^ = 0.23; Experiment 2: *F*_2,70_ = 18.77, *P* < 0.0000001, η^2^ = 0.19), driven by relatively lower accuracy (higher interference) in the high similarity condition (also see **Supplementary Figure 1**). **(C)** On the color memory Post Tests, the mean percentage of responses away from the competitor varied across similarity conditions (main effect of similarity, *P*s < 0.005 for all Days/Experiments). Repulsion effects (> 50% of responses away from competitor) were selectively observed in the high similarity condition (*P*s < 0.05 for all Days/Experiments). Small dots reflect data from individual subjects. Notes: Error bars reflect +/− S.E.M.; *** *P* < 0.001, * *P* < 0.05, ~ *P* < 0.10.

For the Day 1 Post Test, an ANOVA with similarity condition as a factor revealed a robust main effect of similarity on the percentage of responses away from the competitor (*F*_2,44_ = 10.11, *P* = 0.0002, η^2^ = 0.22; Figure 2C). Critically, there was a strong repulsion effect in the high similarity condition (M = 60.80%; SD = 12.00%; *t*_22_ = 4.31, *P* = 0.0003, 95% CI: 5.61 – 15.99, Cohen’s *d* = 1.27), but not in the moderate similarity (M = 48.70%; SD = 9.37%; *t*_22_ = −0.67, *P* = 0.51, 95% CI: −5.36 – 2.75, Cohen’s *d* = −0.20) or low similarity conditions (M = 49.93%; SD = 10.16 %; *t*_22_ = −0.03, *P* = 0.97, 95% CI: −4.46 – 4.32, Cohen’s *d* = −0.01). Thus, when similarity between competing objects was high, color memory for a target object was systematically biased away from the color of the competing object. Follow-up comparisons confirmed that the percentage of away responses was significantly greater in the high similarity condition compared to both the moderate similarity (*t*_22_ = 4.47, *P* = 0.0002, 95% CI = 6.49 – 17.71, Cohen’s *d* = 1.12) and low similarity conditions (*t*_22_ = 3.80, *P* = 0.001, 95% CI: 4.93 – 16.81, Cohen’s *d* = 0.98). The selectivity of the repulsion effect to the high similarity condition is striking when considering that interference errors during the associative memory tests in the training rounds were much more common in the high similarity condition than in the moderate or low similarity conditions (**Supplementary Figure 2**).

Interestingly, the repulsion effect strongly persisted the following day: the main effect of similarity condition was again significant at the Day 2 Post Test (*F*_2,42_ = 9.82, *P* = 0.0003, η^2^ = 0.19; Figure 2C) and there was a selective repulsion effect in the high similarity condition (high similarity: M = 59.77%; SD = 11.17%; *t*_21_ = 4.11, *P* = 0.0005, 95% CI: 4.82 – 14.72, Cohen’s *d* = 1.24; moderate similarity: M = 47.80%; SD = 10.55%, *t*_21_ = −0.98, *P* = 0.34, 95% CI: −6.87 – 2.48, Cohen’s *d* = −0.29; low similarity M = 49.39%; SD = 12.05%, *t*_21_ = −0.24, *P* = 0.82, 95% CI: −5.95 – 4.73, Cohen’s *d* = −0.07).

### Experiment 2

In a replication study (Experiment 2) we used the same procedure as Experiment 1 except for a few minor changes (see Methods for details and rationale). In particular, we reduced the number of color memory tests during the training rounds (by 50%) and eliminated the Day 2 Post Test. Performance across the training rounds is reported in Figure 2A,B and **Supplementary Figures 1** and **2**. As in Experiment 1, the percentage of responses away from the competitor during the color memory Post Test robustly varied across color similarity conditions (*F*_2,70_ = 6.79, *P* = 0.002, η^2^ = 0.09; Figure 2C). Critically, we again observed a significant repulsion effect in the high similarity condition (M = 54.63%; SD = 12.74%; *t*_35_ = 2.16, *P* = 0.036, 95% CI: 0.32 – 8.95, Cohen’s *d* = 0.51). The percentage of away responses did not significantly differ from 50% in the moderate similarity condition (M = 47.95%; SD = 10.42%; *t*_35_ = −1.18, *P* = 0.25, 95% CI: −5.57 – 1.48, Cohen’s *d* = −0.28) and there was a marginally-significant effect in the opposite direction (below 50%) in the low similarity condition (M = 46.05%; SD = 12.07%; *t*_35_ = 1.96, *P* = 0.057, 95% CI: −8.04 – 0.13, Cohen’s *d* = −0.46). Follow-up tests confirmed that the percentage of away responses was again significantly higher in the high similarity condition compared to both the moderate similarity condition (*t*_35_ = 3.10, *P* = 0.004, 95% CI: 2.30 – 11.06, Cohen’s *d* = 0.57) and the low similarity condition (*t*_35_ = 3.36, *P* = 0.002, 95% CI: 3.40 – 13.77, Cohen’s *d* = 0.69). Thus, as in Experiment 1, we observed a selective repulsion effect in color memory specifically when there was high similarity between competing objects.

### Experiment 3

Experiments 1 and 2 strongly establish that the repulsion effect is competition dependent in that it was selective to the high similarity condition. Interestingly, however, while hippocampal repulsion effects are also competition dependent, the relationship between competition and repulsion is thought to be non-monotonic: that is, with sufficiently strong competition, representations will fail to diverge (Ritvo, Turk-Browne, & Norman, 2019). In Experiment 3, we tested whether there was a non-monotonic relationship between color similarity and repulsion by shifting the range of color similarity to include a moderate similarity condition (again, 48 degrees), a high similarity condition (again, 24 degrees) and a new ‘ultra similarity’ condition (6 degrees; Figure 3A).

**Figure 3:**
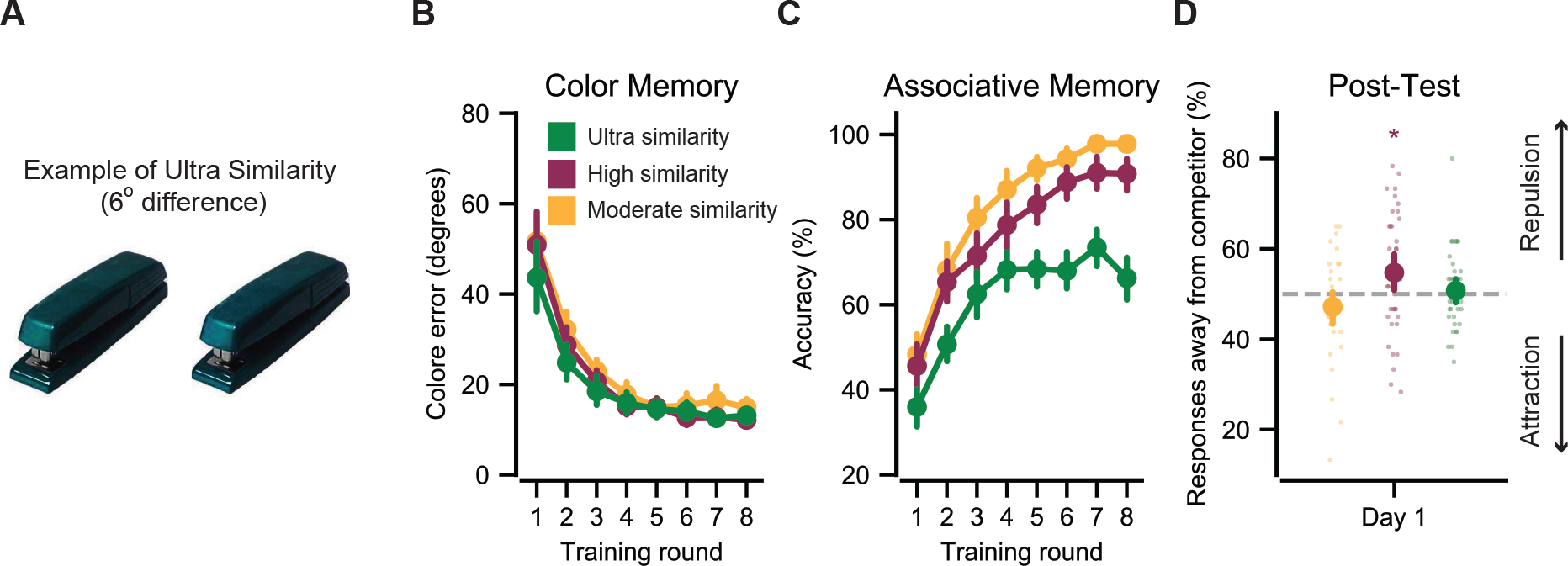
Non-monotonic relationship between color similarity and repulsion. **(A)** Experiment 3 used three color similarity conditions: moderate (48 degrees apart), high (24 degrees), and ultra (6 degrees). An example of competing images from the ultra similarity condition is shown. **(B)** Mean color memory error decreased across training rounds (main effect of round: *F*_1,37_ = 186.5, *P* < 0.0000001, η^2^ = 0.70). **(C)** Accuracy on the associative memory tests increased across training rounds (main effect of test round: *F*_1,37_ = 326.9, *P* < 0.0000001, η^2^ = 0.75). Accuracy differed across color similarity conditions (main effect of similarity: *F*_2,74_ = 129.9, *P* < 0.00000001, η^2^ = 0.60), driven by relatively lower accuracy (higher interference) in the ultra similarity condition (also see **Supplementary Figure 2**). **(D)** On the color memory Post Test, the mean percentage of responses away from the competitor varied across similarity conditions (main effect of similarity: *P =* 0.006). A repulsion effect was selectively observed in the high similarity condition (*P* = 0.036). Small dots reflect data from individual subjects. Notes: Error bars reflect +/− S.E.M.; * *P* < 0.05.

Performance across the training rounds is reported in Figure 3B,C and **Supplementary Figures 1** and **2**. Of particular relevance, associative memory test accuracy (during the training rounds) was significantly lower in the ultra similarity condition than the high similarity condition (*t*_37_ = −11.39, *P* < 0.00000001, 95% CI: −17.95 – −12.53, Cohen’s *d* = −1.85) or moderate similarity condition (*t*_37_ = −16.26, *P* < 0.00000001, 95% CI: −24.26 – −18.86, Cohen’s *d* = −2.96) (Figure 3C and **Supplementary Figure 2**). Nonetheless, in the last associative memory test round, subjects selected the target faces at above-chance rates in all similarity conditions (chance = 25%, all means > 66%, *P*s < 0.0000001). Thus, the ultra similarity condition clearly increased interference relative to the high similarity condition, but subjects were still generally successful at memory-based discrimination between these extremely similar colors.

Results from the Post Test again revealed that color similarity influenced the percentage of responses away from the competitor (*F*_2,74_ = 5.45, *P* = 0.006, η^2^ = 0.06; Figure 3D). However, the relationship between similarity and repulsion followed the predicted non-monotonic pattern. As in Experiments 1 and 2, there was a significant repulsion effect in the high similarity condition (M = 54.74%; SD = 13.42%; *t*_37_ = 2.17, *P* = 0.036, 95% CI: 0.32 – 9.15, Cohen’s *d* = 0.50), no repulsion effect in the moderate similarity condition (M = 47.19%; SD = 10.93%; *t*_37_ = −1.58, *P* = 0.12, 95% CI: −6.40 – 0.79, Cohen’s *d* = −0.36) and a significant difference between the high and moderate similarity conditions (*t*_37_ = 3.04, *P* = 0.004, 95% CI: 2.51 – 12.58, Cohen’s *d* = 0.61). In the ultra similarity condition, however, the percentage of away responses did not differ from 50% (M = 50.75%; SD = 8.52%; *t*_37_ = 0.54, *P* = 0.59, 95% CI: −2.06 – 3.55, Cohen’s *d* = 0.12) confirming that, with sufficiently high similarity, the repulsion effect was eliminated. While the percentage of away responses was numerically lower in the ultra similarity condition than the high similarity condition, this difference did not reach significance (*t*_37_ = −1.63, *P* = 0.11, 95% CI: −0.10 – 8.94, Cohen’s *d* = 0.35). Interestingly, despite the much higher rate of interference errors in the ultra similarity condition compared to the moderate similarity condition (**Supplementary Figure 2**), the percentage of color responses away from the competing object’s color was marginally higher in the ultra similarity condition than in the moderate similarity condition (*t*_37_ = 1.89, *P* = 0.067, CI: −0.026 – 7.37, Cohen’s *d* = 0.36). Taken together, performance across the three similarity conditions suggests a ‘local maximum’ in the repulsion effect that occurred when similarity was high (24 degrees) but not too high (6 degrees).

### Relationship between repulsion and memory interference

Thus far, we have shown that the repulsion effect is *triggered by* similarity between memories. This raises the complementary question: what is the *consequence* of repulsion? From an adaptive perspective, repulsion may carry an important benefit in that, by exaggerating the differences between similar memories, it serves to *reduce* memory interference. To test for a relationship between repulsion and interference, we considered data from Experiments 1–3 and focused specifically on the high similarity condition since a significant repulsion effect was observed in this condition across all three experiments. For each subject in each experiment, we computed (1) the mean percentage of responses away from the competitor based on data from the immediate Post Test and (2) the mean number of interference errors across the last three rounds of the associative memory test (during the training rounds). As a first step, we tested for across-subject correlations between mean percentage of away responses on the color memory Post Test and mean interference errors on the associative memory test. Strikingly, a highly significant, negative correlation was observed for each experiment (Experiment 1: *r* = −0.61, *P* = 0.002; Experiment 2: *r* = −0.51, *P* = 0.001; Experiment 3: *r* = −0.44, *P* = 0.006; Figure 4A,B). Thus, stronger color memory repulsion was associated with fewer interference-related errors during the associative memory test. One potential caveat with the correlations described above is that they may partly reflect that subjects who suffered more interference errors during associative memory tests had a higher probability of remembering the wrong color (i.e., the competing object’s color) during the color memory Post Test. From this perspective, it is possible that all subjects showed comparable levels of repulsion *when they recalled the correct color*, but subjects with more interference errors also recalled the ‘wrong color’ with greater frequency, thereby pulling down their percentage of responses away from the competitor. To address this concern, we performed a second, more targeted analysis that focused on the distribution of *correct* color memory responses. We first median split all subjects (within each experiment) into ‘high interference’ and ‘low interference’ groups based on the mean number of interference errors during the last three associative memory tests in the training rounds (high interference group: M = 18.45%, SD = 7.58%; low interference group: M = 2.12%, SD = 2.59%). We then computed the frequency of Post Test responses that fell in each of four color space bins. Two of these bins were centered around the target color value (+/– 11 degrees from the target color) and two of these bins were centered around the competitor color value (+/– 11 degrees from the competitor color) (Figure 4C). This allowed us to separate out color memory responses that were ‘correct’ (+/– 11 degrees from the target) vs. ‘swap errors’ (+/– 11 degrees from the competitor). Note: responses that were precisely equal to the target or competitor color were excluded from this analysis.

**Figure 4.**
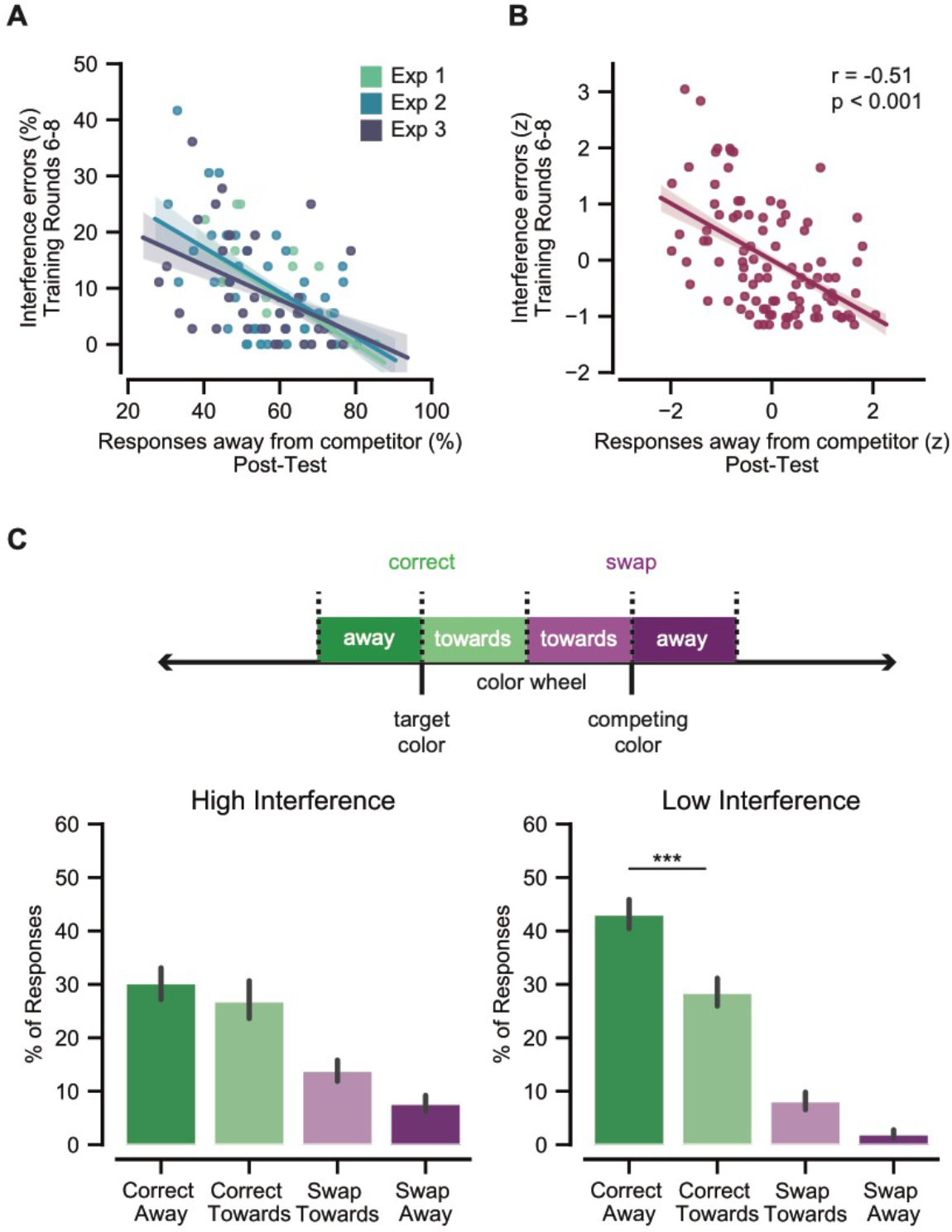
Memory repulsion is adaptive. **(A)** Across-subject correlations between mean percentage of Post Test (Day 1) color memory responses that were away from the competitor and mean percentage of interference errors during the last three associative memory test rounds (high similarity condition only). Interference errors were defined as selecting the face associated with the competitor object (see **Figure 1B** and **Supplementary Figure 2**). Significant, negative correlations were observed for Experiments 1, 2, and 3 (*r*s >. 43, *P*s < 0.007), indicating that stronger color memory repulsion was associated with fewer interference errors during the associative memory test. **(B)** Same as **(A)**, except that each measure was *z*-scored within experiment, allowing for a single correlation to be calculated for the pooled data (*r* = −0.51, *P* < 0.001). **(C)** High interference subjects evenly distributed ‘correct’ color memory responses around the target value (correct toward vs. correct award: *P* = 0.16). In contrast, low interference subjects exhibited a strong bias in their distribution of correct color responses, with significantly more correct responses ‘away’ from the competing color than ‘towards’ the competing color (*P* < 0.0001). The distribution of correct responses (correct towards vs. correct away) significantly interacted with subject group (high vs. low interference; *P* = 0.0006). Notes: Error bars reflect +/− S.E.M.; *** *P* < 0.001.

Not surprisingly, high interference subjects tended to commit more swap errors in the color memory test (*M* = 21.47% of responses, *SD* = 9.51%) than did low interference subjects (*M* = 10.07%, *SD* = 7.28%); (*t*_82_ = 6.17, *P* = < 0.001, 95% CI: 7.73 −15.10, Cohen’s *d* = 1.35). Of critical interest, however, was the distribution of correct responses—i.e., whether there was a difference in the frequency of ‘correct towards’ vs. ‘correct away’ responses. For high interference subjects, correct color memory responses were evenly distributed around the actual target value (no difference in frequency of ‘correct towards’ vs. ‘correct away’: *F*_1,39_ = 2.07, *P* = 0.16, η^2^ = 0.03; left panel Figure 4C). For low interference subjects, however, there was a strong asymmetry, with significantly more responses in the ‘correct away’ bin as compared to the ‘correct towards’ bin (*F*_1,39_ = 58.00, *P* < 0.0001, η^2^ = 0.39; right panel Figure 4C). The interaction between high vs. low interference subjects and ‘correct away’ vs. ‘correct towards’ bins was also highly significant (*F*_1,78_ = 33.33, *P* = 0.0006, η^2^ = 0.08). None of these effects interacted with experiment number (*P*s > 0.4). Thus, even when restricting focus to color memory responses that were correct (i.e., removing swap errors), there was clear evidence for an adaptive distortion of color memory: subjects that made the fewest interference errors during the associative memory test exhibited a robust repulsion effect wherein color memory was systematically biased away from the color of the competing object.

### Experiment 4

In Experiments 1-3, the associative memory task during the training rounds explicitly required subjects to discriminate between the competing objects. In Experiment 4, we tested whether this discrimination demand was necessary for inducing repulsion. The critical difference in Experiment 4, relative to Experiments 1–3, was that we changed the procedures for the associative memory test in the training rounds so that it now promoted *integration* across overlapping associations (Richter, Chanales, & Kuhl, 2016; Shohamy & Wagner, 2008; Zeithamova, Dominick, & Preston, 2012). Specifically, the new associative memory test was an inference test (Zeithamova et al., 2012) that required subjects to generalize across overlapping associations. On each trial in the inference test, a face image (probe) was presented at the top of the screen and subjects had to select a “matching” face, from a set of 4 options presented below (Figure 5A). A “matching” face was defined as a face associated with the same object category as the probe (ignoring differences in color). For example, two faces would match if they were each associated with a “backpack,” regardless of whether the backpacks differed in color. Thus, while the inference test still required associative learning (i.e., object-face learning), it did not require discriminating between similar objects. However, because color memory was still tested during the training rounds (as in all prior experiments), color memory remained relevant and subjects showed robust improvements in color memory across training rounds (Figure 5B).

**Figure 5.**
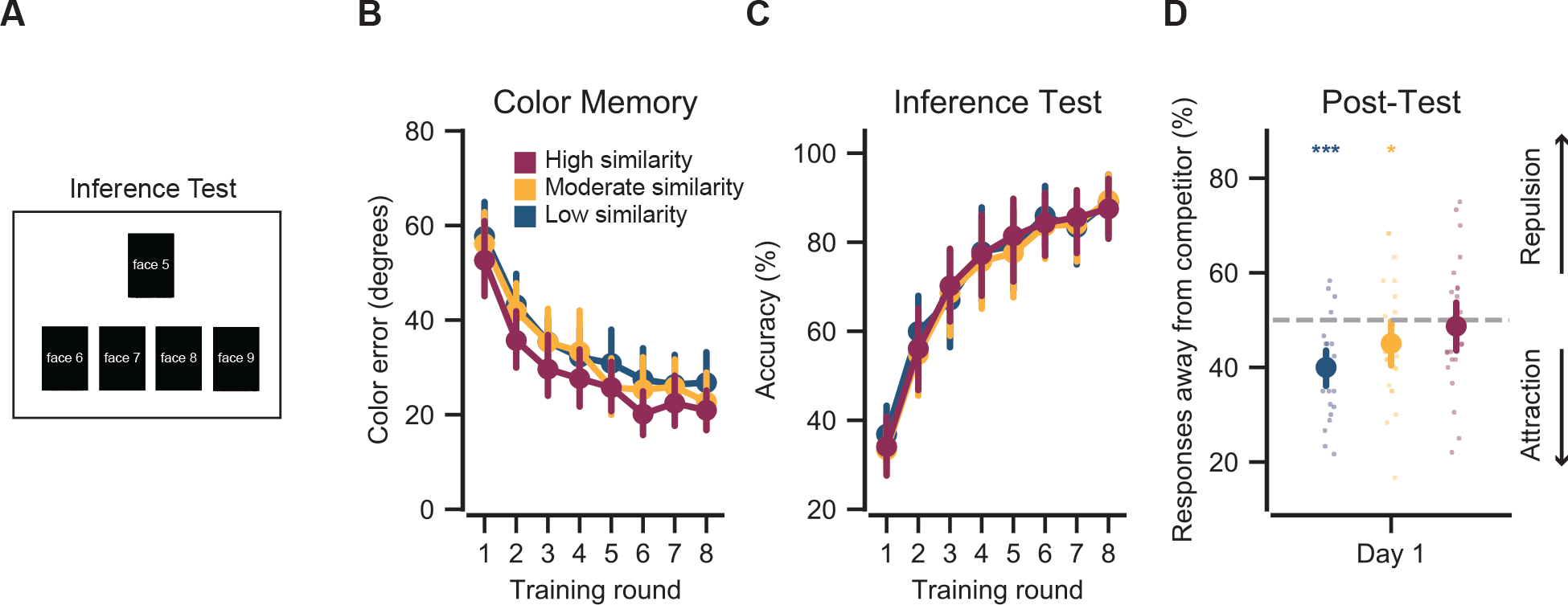
Task demands influence memory repulsion. **(A)** In Experiment 4, all procedures were identical to Experiment 1 except for a subtle change to the associative memory test during the training rounds. Instead of requiring subjects to *discriminate* between competing colors (as in Experiment 1), the associative memory test consisted of an inference test that required subjects to *generalize* across competing objects. On each inference test trial, a probe face was presented and subjects had to select, from a set of four options, which face was associated with the same object as the probe face (irrespective of color). Thus, what was previously the ‘competitor’ face (Experiments 1–3) was now the correct response. Note: Actual faces are not shown here per biorxiv policy. **(B)** Mean color memory error decreased across training rounds (main effect of round: *F*_1,25_ = 101.03, *P* < 0.0000001, η^2^ = 0.63). **(C)** Accuracy on the associative memory tests increased across training rounds (main effect of round: *F*_1,25_ = 225.3, *P* < 0.00000001, η^2^ = 0.79). However, in contrast to the associative memory tests in Experiments 1–3, there was no effect of color similarity on inference accuracy (main effect of similarity: *F*_2,50_ = 0.52, *P* = 0.67, η^2^ = 0.006). **(D)** On the color memory Post Test, the mean percentage of responses away from the competitor varied across similarity conditions, as in Experiments 1–3 (main effect of similarity: *P =* 0.009), but there was no longer a repulsion effect in the high similarity condition (*P* = 0.63). Instead, the mean percentage of responses away from the competitor was significantly below 50% in the moderate and low similarity conditions (*P*s < 0.05). Small dots reflect data from individual subjects. Notes: Error bars reflect +/− S.E.M.; *** *P* < 0.001; * *P* < 0.05.

Subjects performed well in the inference task, with above-chance performance in all conditions by the last round of training (*P*s < 0.000001; Figure 5C). However, in contrast to what we found in the associative memory tests in Experiments 1–3, color similarity had no effect on performance in the inference test (*F*_2,50_ = 0.52, *P* = 0.67, η^2^ = 0.006; Figure 5C). That is, the ability to generalize across associations with common object categories was not influenced by color similarity.

As in all of the prior experiments, the percentage of responses away from the competitor during the color memory Post Test varied by similarity condition (*F*_2,50_ = 5.21, *P* = 0.009, η^2^ = 0.09; Figure 5D). However, for the high similarity condition the percentage of away responses no long differed from 50% (*t*_25_ = −0.49, *P* = 0.63, 95% CI: −6.64 – 4.07, Cohen’s *d* = −0.14). Interestingly, the percentage of away responses was significantly *lower* than 50% in both the moderate and low similarity conditions (moderate: *t*_25_ = −2.08, *P* = 0.048, 95% CI: −9.83 – −0.04, Cohen’s *d* = −0.58; low: *t*_25_ = −5.17, *P* = 0.00002, 95% CI: −13.99 – −6.02, Cohen’s *d* = −1.43), suggesting an attraction effect. A direct comparison of the percentage of away responses in the high similarity conditions in Experiment 4 vs. Experiment 1 (which was most closely matched to Experiment 4) revealed a significant difference (*t*_47_ = 3.33, *P* = 0.002, 95% CI: 4.77 – 19.39, Cohen’s *d* = 0.95), confirming that the subtle change in task demands (encouraging integration as opposed to discrimination) significantly reduced the repulsion effect.

## DISCUSSION

While numerous studies have documented the situations and contexts in which interference between episodic memories produces forgetting, much less is known about how interference shapes memory for the *features* of events. Using a behavioral paradigm that assessed color memory on a continuous scale, we show that interference between similar-colored objects induces a repulsion effect wherein the colors of these objects are remembered as being farther apart than they actually are. This repulsion effect was highly dependent on competition (color similarity) and was sensitive to task demands. Critically, repulsion was also adaptive in that greater repulsion was strongly associated with fewer interference-related errors during associative memory retrieval. These findings provide striking evidence of adaptive memory distortions that are triggered by competition between highly similar memories.

Our study design was modeled after the canonical A-B, A-C interference paradigm (Barnes & Underwood, 1959). In this paradigm, a single memory cue (A) is paired with two different associates (B, C). Memory is typically worse for associations in this interference condition compared to a condition in which a cue is paired with a single associate. In our experiments, however, overlapping associations had *similar cues* (e.g., two backpacks of different colors) as opposed to identical cues (the same backpack). This allowed us to parametrically manipulate the overlap between A-B, A-C associations (conceptually: A_1_-B, A_2_-C). Central to our predictions was the idea that greater similarity of the cues (A_1_, A_2_) should be associated with greater interference (O’Reilly, 2010). Performance from the associative memory tests clearly confirmed this prediction (**Supplementary Figure 2**). However, in contrast to classic memory interference paradigms, overall associative memory accuracy was not our critical dependent measure; rather, we used this paradigm to measure distortions in how the cues (object colors) were remembered. If greater similarity between cues leads to greater interference, then we reasoned that exaggerating the *difference* between the cues would be an adaptive mechanism for reducing interference. This is precisely what we found.

Notably, our core predictions concerning memory repulsion effects were directly inspired by recent evidence of repulsion effects in human hippocampal activity patterns (Ballard et al., 2019; Chanales et al., 2017; Favila et al., 2016; Hulbert & Norman, 2015; Schapiro et al., 2012). Indeed, there are a number of striking parallels between these prior neuroimaging findings and the current behavioral findings. First, a critical finding from prior studies is that hippocampal repulsion is *triggered by* event similarity (Chanales et al., 2017; Favila et al., 2016; Hulbert & Norman, 2015; Schapiro et al., 2012). For example, Chanales et al. (2017) found that hippocampal repulsion was greatest for the segments of spatial routes that were most difficult to discriminate. Similarly, Schapiro et al. (2012) found that hippocampal repulsion selectively occurred for abstract images that had high levels of visual similarity. Here, in Experiments 1-3 we found robust and selective evidence of behavioral memory repulsion when color similarity was high (24 degrees apart). Second, hippocampal repulsion is thought to be a gradual, learning-related process (Chanales et al., 2017; Favila et al., 2016; Hulbert & Norman, 2015; Schlichting et al., 2015). Indeed, during initial stages of learning, there may be attraction between hippocampal representations of similar events, with this attraction only ‘flipping’ to repulsion with extended training (Chanales et al., 2017; Favila et al., 2016; Schlichting et al., 2015). Likewise, repulsion effects in the current study only emerged after extensive training; during initial color memory tests (during the training rounds), subjects’ color memories tended to be biased *toward* the competing object’s color (**Supplementary Figure 1**). Finally, hippocampal repulsion is thought to be adaptive in that it is associated with reduced interference (confusability) between highly similar memories (Colgin et al., 2008; Favila et al., 2016; Hulbert & Norman, 2015). Here, we show that the repulsion effect in color memory was, overwhelmingly, more pronounced in those subjects that suffered the fewest interference errors during the associative memory test. When specifically considering high similarity trials with ‘correct’ color memory (defined as +/− 11 degrees of the target), the difference between subjects with high vs. low rates of interference errors was striking: subjects that had more interference errors had response distributions that were centered on the *veridical color value*; in contrast, subjects with fewer interference errors exhibited a response distribution that was *shifted away from the color of competing object*. Thus, the current behavioral findings strongly parallel previously-described properties of hippocampal repulsion.

In order to induce a repulsion effect in color memory, we deliberately developed a training procedure that involved alternation between study and competitive retrieval (Experiments 1–3). This procedure was inspired by evidence that study/retrieval alternation is very effective in creating distinct representations of overlapping memories (Hulbert & Norman, 2015; Storm, Bjork, & Bjork, 2008) and in inducing differentiation of hippocampal activity patterns (Hulbert & Norman, 2015; Kim, Norman, & Turk-Browne, 2017). These dynamics have also been detailed in computational models that address how episodic memory interference is resolved (Norman, Newman, Detre, & Polyn, 2006; Norman, Newman, & Detre, 2007). More generally, our results build on evidence that competitive remembering triggers active mechanisms that reshape the memory landscape in order to reduce interference (Anderson, 2003; Anderson, Bjork, & Bjork, 1994; Kim et al., 2017; Levy, 2002; Norman et al., 2006, 2007; Wimber, Alink, Charest, Kriegeskorte, & Anderson, 2015).

Across our experiments, we identify several boundary conditions for the repulsion effect. First, as noted above, we consistently observed repulsion in the high similarity condition (24 degrees) but not in the moderate/low similarity conditions (48, 72 degrees). However, with even higher similarity (6 degrees), the repulsion effect was no longer significant. Thus, the relationship between similarity and the repulsion effect followed an inverted u-shape function, suggesting a ‘sweet spot’ at which repulsion occurs. This finding is consistent with theoretical perspectives on the relationship between memory competition and memory plasticity (Ritvo et al., 2019). Specifically, memory representations are thought to be most susceptible to plasticity (weakening or distortion) at particular levels of competition. If memory representations are *too similar* or *too dissimilar*, then plasticity is not expected to occur. This theoretical perspective is supported by several examples of non-monotonic relationships between neural measures of competition and memory/plasticity (Chanales et al., 2017; Detre, Natarajan, Gershman, & Norman, 2013; Lewis-Peacock & Norman, 2014; Newman & Norman, 2010).

Another boundary condition to the repulsion effect that we identify relates to task demands. Namely, the repulsion effect was not observed when task demands explicitly encouraged integration (instead of discrimination) of similar objects (Experiment 4). Interestingly, this integration demand led to an ‘attraction effect’ for the low and moderate similarity conditions, but not for the high similarity condition. On the one hand, storing a single ‘averaged’ color value for each object pair (i.e., attraction) would seemingly be an efficient strategy when task demands require integration (Gluck & Myers, 1993; Richards et al., 2014). However, it is possible— though speculative—that event similarity triggers some degree of repulsion regardless of task demands (Favila et al., 2016) and that, in Experiment 4, results in the high similarity condition reflect offsetting effects of integration and repulsion. While detailed consideration of this point is beyond the scope of the present manuscript, our findings establish that task demands are an important factor, along with event similarity, but additional research will be required to map out exactly how and when repulsion effects are influenced by task demands.

Although our findings were specifically motivated by empirical phenomena and theoretical perspectives in the field of episodic memory, they contribute to a broader literature documenting adaptive exaggeration in cognitive processes. For example, similar biases have previously been documented in visual working memory (Bae & Luck, 2017; Rademaker, Bloem, De Weerd, & Sack, 2015), estimates of temporal duration (Ezzyat & Davachi, 2014), and judgments of social categories (Förster, Liberman, & Kuschel, 2008; Krueger & Rothbart, 1990; Wilder & Thompson, 1988). This raises the question of whether the repulsion effect we observed is, fundamentally, a bias in episodic memory or whether the bias might occur during another cognitive processing stage. In particular, it is possible that the bias occurred during perception and this bias was then reinstated during memory retrieval. This framing is not incompatible with our claims. That said, it is important to emphasize that any bias during perception would still be dependent on long-term memory in that a perceptual bias could only occur to the extent that a *remembered stimulus* exerted an influence on a currently perceived stimulus (Teng & Kravitz, 2019). Moreover, it is interesting to note that damage to the hippocampus (a structure critical for episodic memory formation) is also associated with impairments in fine-grained perceptual discriminations (Aly, Ranganath, & Yonelinas, 2013), suggesting that the distinction between memory and perception may not be categorical (Aly & Turk-Browne, 2018). Ultimately, while it is an interesting question whether the repulsion effect reported here also occurred during perception, the critical points are that the repulsion effect we report (a) was induced by long-term memory, (b) it was remarkably stable over time (e.g., it persisted ~24 hours in Experiment 1), and (c) it strongly predicted associative interference errors in a canonical episodic memory paradigm.

Collectively, our results robustly establish that similarity between long-term memories triggers a repulsion in remembered feature values and that this exaggeration of remembered features is highly adaptive. These findings strongly support the idea that memory distortions generally reflect the operation of an adaptive memory system (Schacter, 1999), while providing specific, new evidence of how such distortions can mitigate memory interference.

## Supporting information

Suppelment

## ACKNOWLEDGEMENTS

This work was supported by R01-NS089729 from the National Institute of Neurological Disorders and Stroke to B.A.K.

## REFERENCES

Aly, M., Ranganath, C., & Yonelinas, A. P. (2013). Detecting Changes in Scenes: The Hippocampus Is Critical for Strength-Based Perception. Neuron, 78(6), 1127–1137. https://doi.org/10.1016/j.neuron.2013.04.018

Aly, M., & Turk-Browne, N. B. (2018). Flexible weighting of diverse inputs makes hippocampal function malleable. Neuroscience Letters, 680, 13–22. https://doi.org/10.1016/j.neulet.2017.05.063

Anderson, M. C. (2003). Rethinking interference theory: Executive control and the mechanisms of forgetting. Journal of Memory and Language, 49(4), 415–445. https://doi.org/10.1016/j.jml.2003.08.006

Anderson, M. C., Bjork, R. A., & Bjork, E. L. (1994). Remembering Can Cause Forgetting: Retrieval Dynamics in Long-Term Memory. 25.

Bae, G.-Y., & Luck, S. J. (2017). Interactions between visual working memory representations. Attention, Perception, & Psychophysics, 79(8), 2376–2395. https://doi.org/10.3758/s13414-017-1404-8

Bakker, A., Kirwan, C. B., Miller, M., & Stark, C. E. L. (2008). Pattern Separation in the Human Hippocampal CA3 and Dentate Gyrus. Science, 319(5870), 1640–1642. https://doi.org/10.1126/science.1152882

Ballard, I. C., Wagner, A. D., & McClure, S. M. (2019). Hippocampal pattern separation supports reinforcement learning. Nature Communications, 10(1), 1073. https://doi.org/10.1038/s41467-019-08998-1

Barnes, J. M., & Underwood, B. J. (1959). “Fate” of first-list associations in transfer theory. Journal of Experimental Psychology, 58(2), 97–105. https://doi.org/10.1037/h0047507

Brady, T. F., Konkle, T., Alvarez, G. A., & Oliva, A. (2013). Real-world objects are not represented as bound units: Independent forgetting of different object details from visual memory. Journal of Experimental Psychology: General, 142(3), 791–808. https://doi.org/10.1037/a0029649

Chanales, A. J. H., Oza, A., Favila, S. E., & Kuhl, B. A. (2017). Overlap among Spatial Memories Triggers Repulsion of Hippocampal Representations. Current Biology, 27(15), 2307–2317.e5. https://doi.org/10.1016/j.cub.2017.06.057

Colgin, L. L., Moser, E. I., & Moser, M.-B. (2008). Understanding memory through hippocampal remapping. Trends in Neurosciences, 31(9), 469–477. https://doi.org/10.1016/j.tins.2008.06.008

Detre, G. J., Natarajan, A., Gershman, S. J., & Norman, K. A. (2013). Moderate levels of activation lead to forgetting in the think/no-think paradigm. Neuropsychologia, 51(12), 2371–2388. https://doi.org/10.1016/j.neuropsychologia.2013.02.017

Dimsdale-Zucker, H. R., Ritchey, M., Ekstrom, A. D., Yonelinas, A. P., & Ranganath, C. (2018). CA1 and CA3 differentially support spontaneous retrieval of episodic contexts within human hippocampal subfields. Nature Communications, 9(1), 294. https://doi.org/10.1038/s41467-017-02752-1

Ezzyat, Y., & Davachi, L. (2014). Similarity Breeds Proximity: Pattern Similarity within and across Contexts Is Related to Later Mnemonic Judgments of Temporal Proximity. Neuron, 81(5), 1179–1189. https://doi.org/10.1016/j.neuron.2014.01.042

Favila, S. E., Chanales, A. J. H., & Kuhl, B. A. (2016). Experience-dependent hippocampal pattern differentiation prevents interference during subsequent learning. Nature Communications, 7(1), 11066. https://doi.org/10.1038/ncomms11066

Förster, J., Liberman, N., & Kuschel, S. (2008). The effect of global versus local processing styles on assimilation versus contrast in social judgment. Journal of Personality and Social Psychology, 94(4), 579–599. https://doi.org/10.1037/0022-3514.94.4.579

Gluck, M. A., & Myers, C. E. (1993). Hippocampal Mediation of Stimulus Representation: A Computational Theory. Hippocampus, 3(4), 491–516.

Hulbert, J. C., & Norman, K. A. (2015). Neural Differentiation Tracks Improved Recall of Competing Memories Following Interleaved Study and Retrieval Practice. Cerebral Cortex, 25(10), 3994–4008. https://doi.org/10.1093/cercor/bhu284

Kim, G., Norman, K. A., & Turk-Browne, N. B. (2017). Neural Differentiation of Incorrectly Predicted Memories. The Journal of Neuroscience, 37(8), 2022–2031. https://doi.org/10.1523/JNEUROSCI.3272-16.2017

Krueger, J., & Rothbart, M. (1990). Contrast and Accentuation Effects in Category Learning. Journal of Personality and Social Psychology, 59(4), 651–663.

Levy, B. (2002). Inhibitory processes and the control of memory retrieval. Trends in Cognitive Sciences, 6(7), 299–305. https://doi.org/10.1016/S1364-6613(02)01923-X

Lewis-Peacock, J. A., & Norman, K. A. (2014). Competition between items in working memory leads to forgetting. Nature Communications, 5(1), 5768. https://doi.org/10.1038/ncomms6768

Newman, E. L., & Norman, K. A. (2010). Moderate Excitation Leads to Weakening of Perceptual Representations. Cerebral Cortex, 20(11), 2760–2770. https://doi.org/10.1093/cercor/bhq021

Norman, K. A., Newman, E., Detre, G., & Polyn, S. (2006). How Inhibitory Oscillations Can Train Neural Networks and Punish Competitors. Neural Computation, 18(7), 1577–1610. https://doi.org/10.1162/neco.2006.18.7.1577

Norman, K. A., Newman, E. L., & Detre, G. (2007). A neural network model of retrieval-induced forgetting. Psychological Review, 114(4), 887–953. https://doi.org/10.1037/0033-295X.114.4.887

O’Reilly, R. C. (2010). The Division of Labor Between the Neocortex and Hippocampus. In Connectionist Modeling in Cognitive (Neuroscience) (p. 17). Psychology Press.

Rademaker, R. L., Bloem, I. M., De Weerd, P., & Sack, A. T. (2015). The impact of interference on short-term memory for visual orientation. Journal of Experimental Psychology: Human Perception and Performance, 41(6), 1650–1665. https://doi.org/10.1037/xhp0000110

Richards, B. A., Xia, F., Santoro, A., Husse, J., Woodin, M. A., Josselyn, S. A., & Frankland, P. W. (2014). Patterns across multiple memories are identified over time. Nature Neuroscience, 17(7), 981–986. https://doi.org/10.1038/nn.3736

Richter, F. R., Chanales, A. J. H., & Kuhl, B. A. (2016). Predicting the integration of overlapping memories by decoding mnemonic processing states during learning. NeuroImage, 124, 323–335. https://doi.org/10.1016/j.neuroimage.2015.08.051

Ritvo, V. J. H., Turk-Browne, N. B., & Norman, K. A. (2019). Nonmonotonic Plasticity: How Memory Retrieval Drives Learning. Trends in Cognitive Sciences, 23(9), 726–742. https://doi.org/10.1016/j.tics.2019.06.007

Schacter, D. L. (1999). The seven sins of memory: Insights from psychology and cognitive neuroscience. American Psychologist, 54(3), 182–203.

Schapiro, A. C., Kustner, L. V., & Turk-Browne, N. B. (2012). Shaping of Object Representations in the Human Medial Temporal Lobe Based on Temporal Regularities. Current Biology, 22(17), 1622–1627. https://doi.org/10.1016/j.cub.2012.06.056

Schlichting, M. L., Mumford, J. A., & Preston, A. R. (2015). Learning-related representational changes reveal dissociable integration and separation signatures in the hippocampus and prefrontal cortex. Nature Communications, 6(1), 8151. https://doi.org/10.1038/ncomms9151

Shohamy, D., & Wagner, A. D. (2008). Integrating Memories in the Human Brain: Hippocampal-Midbrain Encoding of Overlapping Events. Neuron, 60(2), 378–389. https://doi.org/10.1016/j.neuron.2008.09.023

Storm, B. C., Bjork, E. L., & Bjork, R. A. (2008). Accelerated relearning after retrieval-induced forgetting: The benefit of being forgotten. Journal of Experimental Psychology: Learning, Memory, and Cognition, 34(1), 230–236. https://doi.org/10.1037/0278-7393.34.1.230

Teng, C., & Kravitz, D. J. (2019). Visual working memory directly alters perception. Nature Human Behaviour, 3(8), 827–836. https://doi.org/10.1038/s41562-019-0640-4

Wilder, D. A., & Thompson, J. E. (1988). Assimilation and Contrast Effects in the Judgments of Groups. Journal of Personality and Social Psychology, 54(1), 62–73.

Wimber, M., Alink, A., Charest, I., Kriegeskorte, N., & Anderson, M. C. (2015). Retrieval induces adaptive forgetting of competing memories via cortical pattern suppression. Nature Neuroscience, 18(4), 582–589. https://doi.org/10.1038/nn.3973

Yassa, M. A., & Stark, C. E. L. (2011). Pattern separation in the hippocampus. Trends in Neurosciences, 34(10), 515–525. https://doi.org/10.1016/j.tins.2011.06.006

Zeithamova, D., Dominick, A. L., & Preston, A. R. (2012). Hippocampal and Ventral Medial Prefrontal Activation during Retrieval-Mediated Learning Supports Novel Inference. Neuron, 75(1), 168–179. https://doi.org/10.1016/j.neuron.2012.05.010

