## Supplementary material for "Adaptive repulsion of long-term memory representations is triggered by event similarity": Suppelment

### SUPPLEMENTARY FIGURES

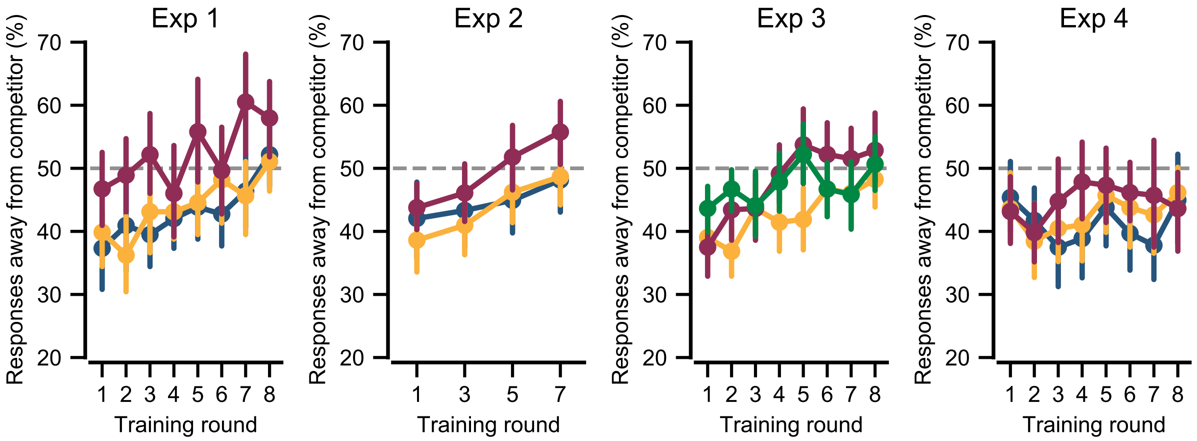


**Supplementary Figure 1**. Mean percentage of responses away from the competitor during the color memory tests, as a function of similarity condition and training round, for Experiments 1–4. In Experiments 1 and 2, the percentage of responses away from the competitor increased across training rounds (main effect of training round, Experiment 1: $F_{1,22}$ = 20.34, $P$ = 0.0002; Experiment 2: $F_{1,34}$ = 23.66, $P$ = 0.00003) and differed as a function of color similarity (Experiment 1: $F_{2,44}$ = 7.78, $P$ = 0.001; Experiment 2: $F_{2,68}$ = 3.98, $P$ = 0.023), with relatively more responses away from the competitor in the high similarity condition. In Experiment 3, the percentage of responses away from the competitor increased across training rounds (main effect of training round: $F_{1,37}$ = 30.92, $P$ = 0.000002) and differed as a function of color similarity ($F_{2,74}$ = 4.85, $P$ = 0.01), with relatively more responses away from the competitor in the ultra and high similarity conditions. In Experiment 4, the percentage of responses away from the competitor did not increase across training rounds (main effect of training round: $F_{1,25}$ = 0.85, $P$ = 0.37) or differ as a function of color similarity ($F_{2,50}$ = 1.38, $P$ = 0.27).

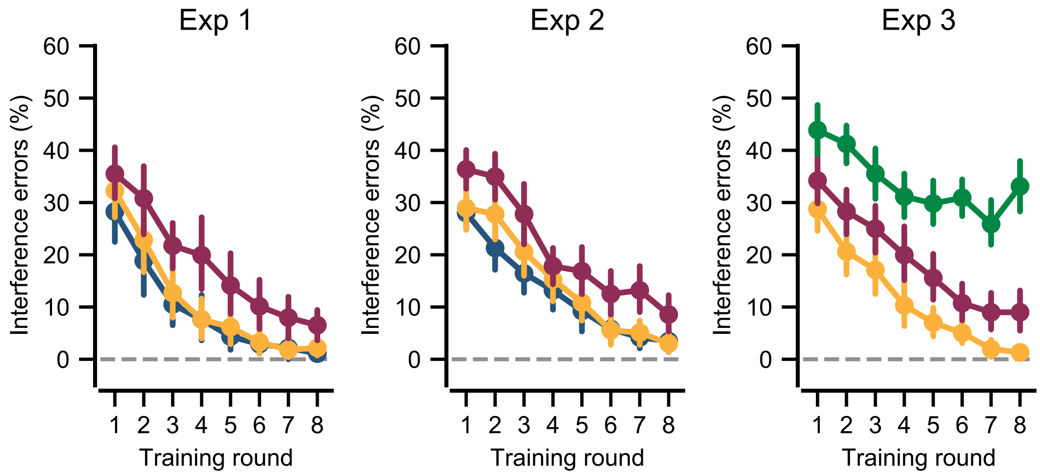


**Supplementary Figure 2**. Mean percentage of interference errors during the associative memory tests, as a function of similarity condition and training round, for Experiments 1–3. Interference errors occurred when subjects selected the face that had been associated with the competitor object. In Experiment 1, there were significant main effects of training round ($F_{1,22}$ = 230.4, $P$ < 0.001) and color similarity ($F_{2,44}$ = 20.49, $P$ < 0.001), with relatively more interference errors in the high similarity condition. In Experiment 2, there were significant main effects of training round ($F_{1,35}$ = 489.0, $P$ < 0.001) and color similarity ($F_{2,70}$ = 24.43, $P$ < 0.001), with relatively more interference errors in the high similarity condition. In Experiment 3, there were significant main effects of training round ($F_{1,37}$ = 188.1, $P$ < 0.001) and color similarity ($F_{2,74}$ = 163.5 $P$ < 0.001), with relatively more interference errors in the ultra similarity condition than high similarity condition and relatively more interference errors in the high similarity condition than the moderate similarity condition.

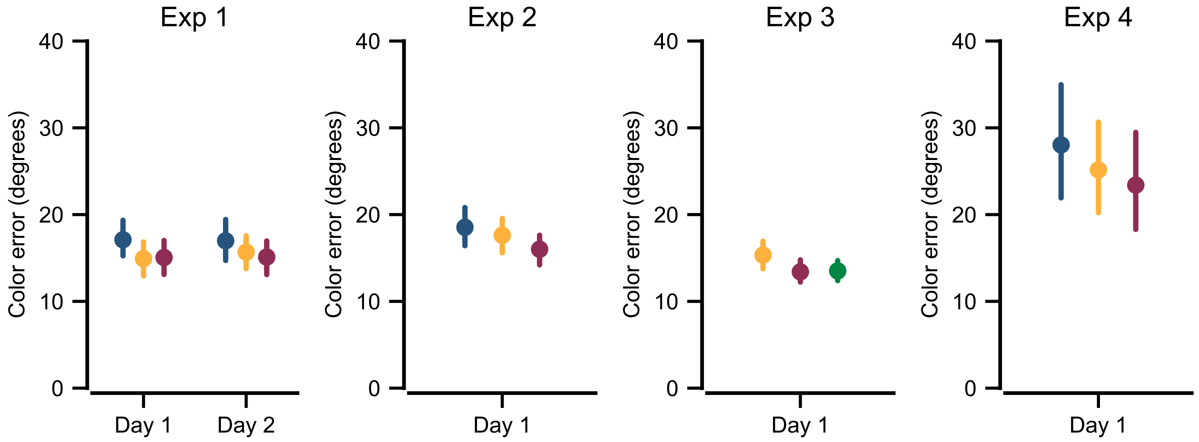


**Supplementary Figure 3**. Mean error of Post Test color memory responses, as a function of similarity condition, for Experiments 1–4. For Experiment 1, color error varied across color similarity conditions on Day 1 ($F_{2,44}$ = 4.03, $P$ = 0.025), with a similar trend on Day 2 ($F_{2,42}$ = 2.35, $P$ = 0.11). For Experiment 2, color error varied across color similarity conditions ($F_{2,70}$ = 4.99, $P$ = 0.009). In each case, error was relatively greatest in the low similarity condition. For Experiment 3, color error again varied across similarity conditions ($F_{2,74}$ = 6.05, $P$ = 0.004), with the relatively greatest error in the moderate similarity condition (color error did not differ between the high and ultra similarity conditions: $t_{37}$ = -0.17, $P$ = 0.86). For Experiment 4, color error was overall much greater than in any of the prior experiments ($P$s < .05), consistent with the fact that the inference test that was used in Experiment 4 did not require that subjects discriminate between similar colors. Nonetheless, color error varied by similarity condition ($F_{2,50}$ = 6.31, $P$ = 0.004) with relatively greatest color error in the low similarity condition (as in Experiments 1 and 2).
